# The OceanDNA MAG catalog contains over 50,000 prokaryotic genomes originated from various marine environments

**DOI:** 10.1101/2021.08.18.456858

**Authors:** Yosuke Nishimura, Susumu Yoshizawa

## Abstract

Marine microorganisms are immensely diverse and play fundamental roles in global geochemical cycling. Recent metagenome-assembled genome studies, with special attention to large-scale projects such as *Tara* Oceans, have expanded the genomic repertoire of marine microorganisms. However, published marine metagenome data has not been fully explored yet. Here, we collected 2,057 marine metagenomes (>29 Tera bps of sequences) covering various marine environments and developed a new genome reconstruction pipeline. We reconstructed 52,325 qualified genomes composed of 8,466 prokaryotic species-level clusters spanning 59 phyla, including genomes from deep-sea deeper than 1,000 m (n=3,337), low-oxygen zones of <90 μmol O_2_ per kg water (n=7,884), and polar regions (n=7,752). Novelty evaluation using a genome taxonomy database shows that 6,256 species (73.9%) are novel and include genomes of high taxonomic novelty such as new class candidates. These genomes collectively expanded the known phylogenetic diversity of marine prokaryotes by 34.2% and the species representatives cover 26.5 - 42.0% of prokaryote-enriched metagenomes. This genome resource, thoroughly leveraging accumulated metagenomic data, illuminates uncharacterized marine microbial ‘dark matter’ lineages.

## Background & Summary

Marine microorganisms have shaped Earth’s environment and played crucial roles in controlling the global climate^1,2^. Genome-based knowledge is essential to understand microorganisms in various aspects, such as their phylogeny, evolution, metabolism, and physiology. Though difficulty in isolation has limited the genome-based knowledge of marine microorganisms, success of culture independent genome reconstruction techniques such as metagenome-assembled genomes (MAGs) and single-amplified genomes (SAGs) have changed our understanding of microbial ecosystems. Genome information of marine microorganisms supplied by these approaches enabled to uncover new lineages that have been identified as participants in important biogeochemical cycling (e.g., nitrogen fixation^3^ and carbon fixation^4,5^), characterize metabolic potentials of uncultured lineages^6,7,8,9,10^, and reconstruct deep evolutionary trajectories^11,12^.

Metagenomes of *Tara* Oceans Expeditions^13,14^ have been repeatedly subjected for genome reconstruction^3,4,10,11,15,16,17^. In contrast, there are many metagenomes from which relatively little effort has been made for genome reconstruction despite large-scale data (e.g., GEOTRACES^18^) or from which reported genomes were limited to ones of specific taxa (e.g., metagenomes of the Canada Basin^19^). Moreover, genome reconstruction methodologies in many previous studies are considered inefficient (e.g., use of a single binning algorithm and/or coverage profile calculated by a single or only limited samples^20^). Genome reconstruction using an improved methodology and applying it to a large-scale metagenome dataset is thus promising for expanding our genomic knowledge of marine microorganisms.

We aimed to build an extended genome catalog of marine prokaryotes with taking advantage of accumulated metagenomic data. Practically, two methodological focuses of this study were defined as (1) to compose a large-scale metagenome dataset that covers diverse marine environments including less explored regions such as deep-sea, low-oxygen zones, and polar regions and (2) to develop a new genome reconstruction pipeline to maximize quality of reconstructed genomes. Here, we collected 2,057 published metagenomes originated from diverse marine environments (Fig. 1ab). Then, to improve the quality of genomes, we developed a genome reconstruction pipeline that includes three key processes (Fig. 1c). As a result, we reconstructed 52,325 qualified prokaryotic genomes that were QS (quality score: %-completeness - 5 x %-contamination) ≥ 50, named as the OceanDNA MAGs. These genomes were reconstructed from various marine environments, including genomes originated from deep-sea deeper than 1,000 m (n=3,337; from 179 metagenomes), low-oxygen zones of <90 μmol O_2_ per kg water (n=7,884; from 176 metagenomes), and polar regions (n=7,752; from 129 metagenomes) (Fig. 2a).

**Figure 1.**
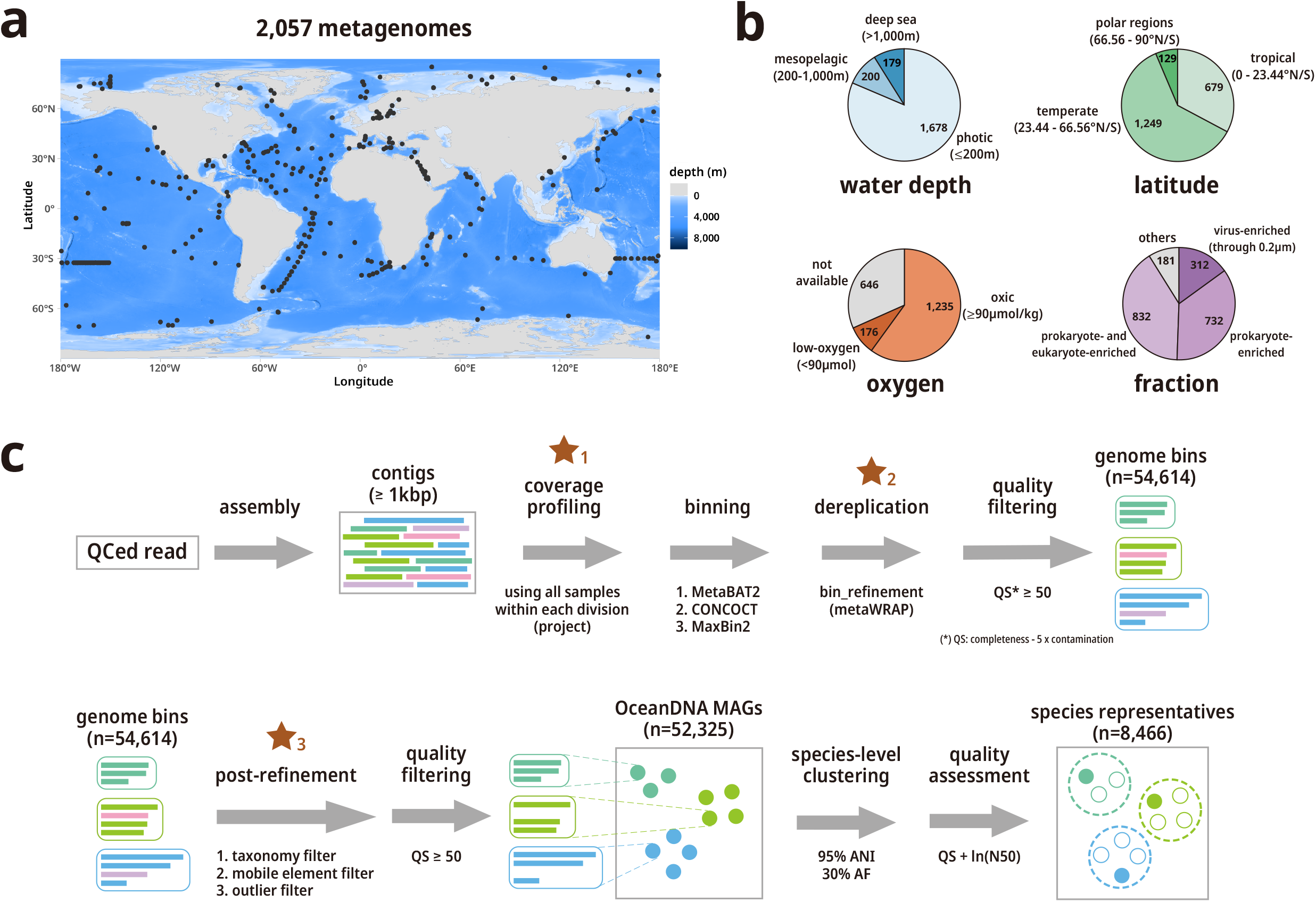
Overview of the study. (a) Geographic distribution of the 2,057 metagenomes analysed in this study (shown by black points). The map was drawn using marmap^70^ and ggplot2 (https://ggplot2.tidyverse.org/). (b) Origin of the metagenome samples. Types of the fraction were described in the main text. (c) Schematic representation of the developed pipeline for MAG reconstruction. Three key processes were highlighted by brown stars. Source data of (a) and (b) was provided in Supplementary File S1.

**Figure 2.**
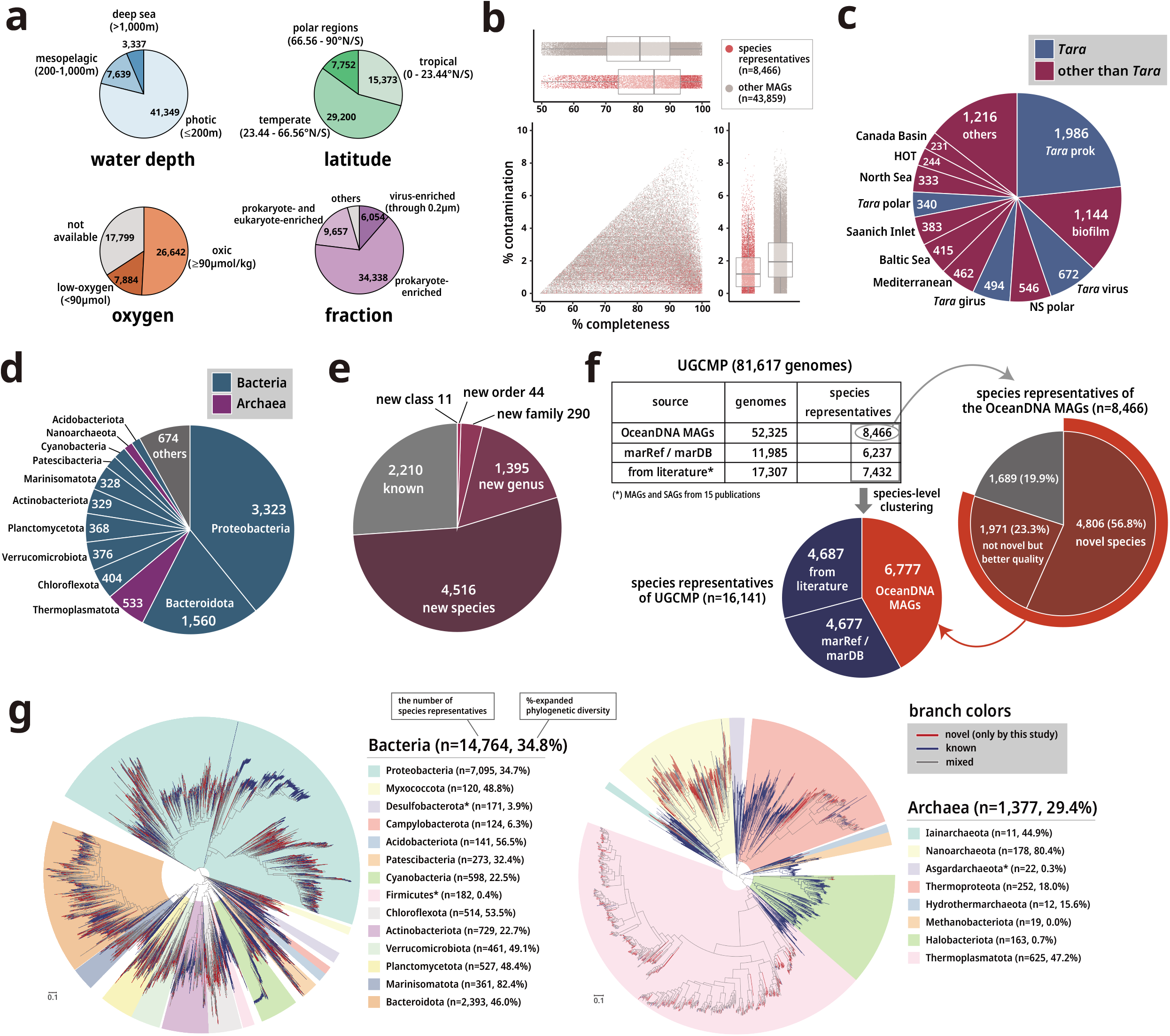
Origin, quality, and novelty of the OceanDNA MAGs. (a) Origin of the OceanDNA MAGs. Types of the fraction were described in the main text. (b) Genome completeness and contamination evaluated by CheckM. (c) Origin of metagenome divisions of the 8,466 species representatives. (d) Phyla of the species representatives assigned by GTDB-Tk. (e) Potential novelty of the species representatives assessed using GTDB-Tk. (f) Origins and compositions of the unified catalog UGCMP and the species representatives. (g) Bacterial (left) and archaeal (right) phylogenetic trees of the species representatives of UGCMP. The trees were midpoint rooted for visualization purpose. The number of species representatives and %-expanded phylogenetic diversity were described for individual phyla of which the number of species was at least 100 for bacteria and 10 for archaea. These phyla were highlighted in the trees with the corresponding colours. If a phylum was not monophyletic in the trees, only the largest monophyletic unit was highlighted (three phyla represented by asterisks in the legend). Note that %-expanded phylogenetic diversity was estimated using all the genomes of UGCMP (not limited to the species representatives).

The OceanDNA MAGs were composed of 8,466 species-level clusters. Genomes were identified as species representatives if the genome quality is the best within each species-cluster (assessed by ‘QS + ln(N50)’). The median genome completeness and contamination of the OceanDNA MAGs were estimated as >80% and <2%, respectively (Fig 2b). The species representatives were originated from various metagenomic projects (divisions), and not dominated by ones from *Tara* Oceans (Fig. 2c). Taxonomic classification based on the genome taxonomy database (GTDB) release 05-RS95^21^ showed that the OceanDNA MAGs covered various marine prokaryotic lineages spanning 59 phyla (Fig. 2d). As taxonomic novelty assessment according to GTDB, 11 species representatives were not assigned to any existing class, suggesting that these species potentially belong to new classes. Likewise, 44 species representatives were suggested to belong to new orders, 290 were to new families, and 1,395 were to new genera (Fig. 2e). Overall, A large part of representatives (n=6,256; 73.9%) was not assigned to existing species in the database.

Novelty of the OceanDNA MAGs were further evaluated by a collection of published marine prokaryotic genomes (n=29,292; QS ≥ 50). Among the 8,466 species representatives, a large part (80.1%) was not overlapped with the published genomes at species level (56.8%) or was overlapped but of superior genome quality (here assessed by ‘QS + ln(N50)’) to the published genomes (23.3%) (Fig. 2f). The OceanDNA MAGs expanded the known phylogenetic diversity of marine prokaryotes by 34.2% (34.8% for bacteria and 29.4% for archaea) that was evaluated by the sum of branch length of bacterial and archaeal phylogenomic trees (Fig. 2g). The species representative genomes collectively covered 26.5 - 42.0% of metagenomic reads of prokaryote-enriched metagenomes at ≥95% nucleotide identity (Fig 3a). The OceanDNA MAG catalog is available as an unprecedented-scale genome resource of marine prokaryotes that enable to characterize microbial ‘dark matter’ lineages and to elucidate yet unsolved questions of marine microbial ecosystems.

**Figure 3.**
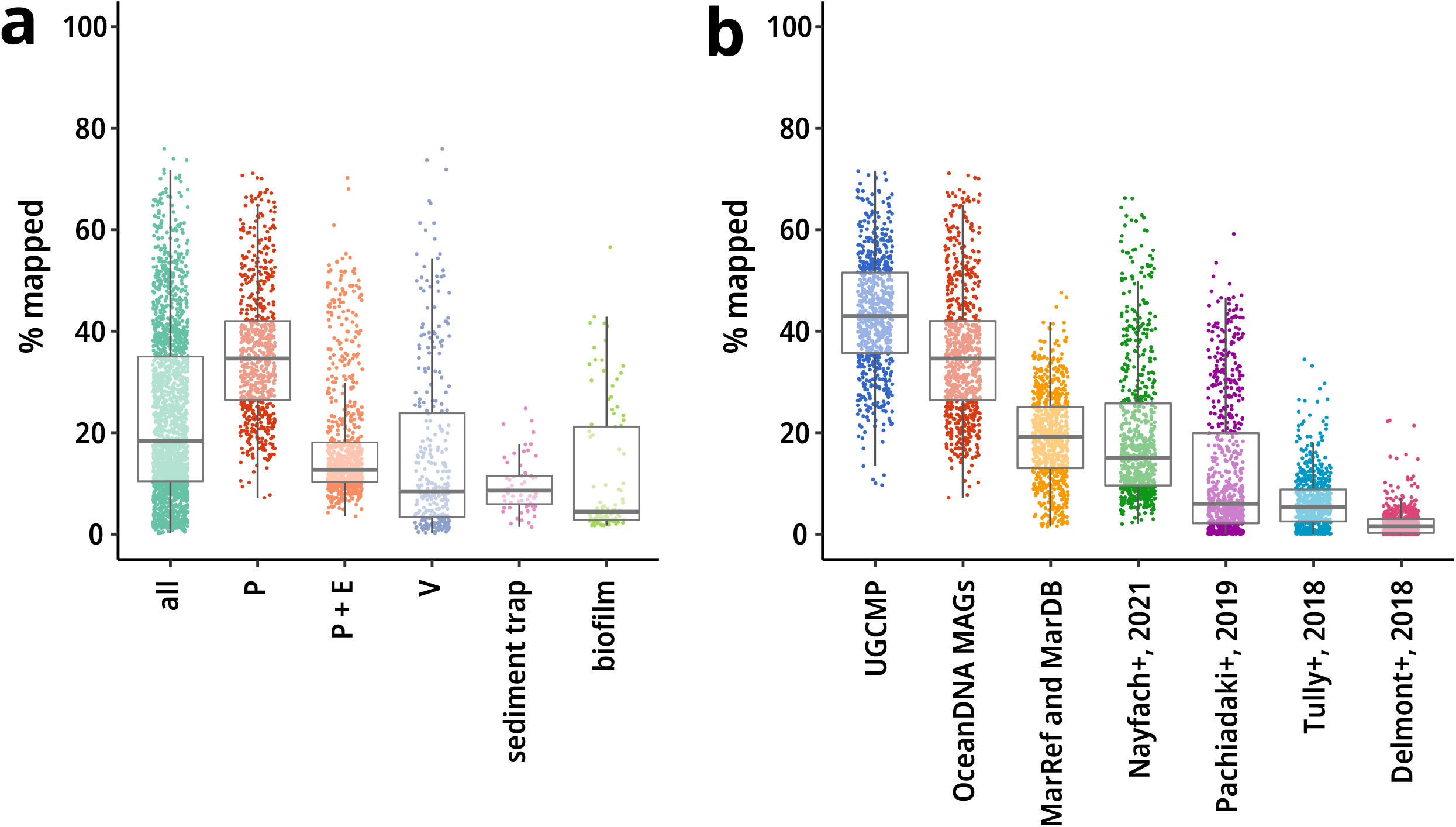
Recruitment of metagenomic reads. The fraction of mapped reads of 2,057 metagenomes were evaluated at ≥95% nucleotide identity. (a) Recruitment onto the species representatives of the OceanDNA MAGs. X-axis shows types of metagenome fractions. P: prokaryote-enriched metagenomes, P + E: prokaryote- and eukaryote-enriched metagenomes, V: virus enriched metagenomes. (b) Recruitment of prokaryote-enriched metagenome reads. X-axis shows genome collections. Note that all these genome collections include only species representatives of qualified genomes (i.e., QS ≥ 50). UGCMP and OceanDNA MAGs include genomes reconstructed in this study. Nayfach+, 2021^62^, Pachiadaki+, 2019^5^, Tully+, 2018^16^, and Delmont+, 2018^3^ are reported genome collections. For Nayfach+, 2021, genomes are limited to ones of which ‘ecosystem type’ is marine.

## Methods

### Collection of metagenomes

We composed a dataset of marine metagenomic samples originated from a broad range of geographic regions (Fig 1ab). These metagenomes were reported by various research groups, and we organized these into 24 divisions for operational purpose, considering various factors such as related publications, research groups, and geographic regions (Table 1). These metagenomes include ones originated from long-distance cruises (e.g., *Tara* Oceans^22,23,24^, GEOTRACES^18^, and Malaspina^25^) and from time-series and/or transect sampling in a specific marine region (e.g., the Mediterranean Sea^26,27^, the Baltic Sea^28^, the Saanich Inlet^29^, Station ALOHA^30^, and the San Pedro Channel^31^). Associated metadata such as location, date, depth, oxygen concentration was collected from original publication and the BioSample database (Supplementary File S1). The metagenomic samples were originated from pole-to-pole (76.96°S - 85.02°N), sea surface to deep-sea (0 - 10,899 m below sea level), oxic to anoxic zones, coastal to pelagic seas (Fig. 1ab). The samples contain ones from aphotic zones (179 metagenomes from deeper than 1,000 m; 200 metagenomes from 200 - 1,000 m) and low-oxygen zones (73 dysoxic (20 - 90 μmol/kg), 86 suboxic (1 - 20 μmol/kg), and 17 anoxic (<1 μmol/kg) metagenomes^32^; Fig 1b). Most samples were originated from prokaryote-enriched fractions (here defined as sea water pass through a prefilter of 0.45 - 5 μm pore and collected on a filter of 0.1 - 0.45 μm pore; n=732), prokaryote- and eukaryote-enriched fractions (pass through a prefilter of 20 μm pore or no prefilter and collected on a filter of 0.2 - 0.8 μm pore; n=832), and virus-enriched fractions (pass through a prefilter of 0.2 - 0.22 μm pore; n=312; Fig 1b). In addition to water samples, metagenomes originated from sediment traps^33, 34^ (n=63) and in situ formation of biofilms^35^ (n=104) were collected. Overall, these metagenomes cover various marine environments.

**Table 1.** metagenome divisions.

| division name | related publication (selected) | samples | QCed read (Gbp) | MAGs |
| --- | --- | --- | --- | --- |
| *Tara* prok | Sunagawa et al., 2015<sup>22</sup> | 139 | 4,935 | 8,624 |
| Saanich Inlet | Hawley et al., 2017<sup>29</sup> | 85 | 1,041 | 5,087 |
| NS polar | Cao et al., 2020<sup>58</sup> | 59 | 847 | 3,511 |
| *Tara* virus | Gregory et al., 2019<sup>23</sup> | 131 | 3,887 | 3,271 |
| Monterey bloom | Nowinski et al., 2019<sup>44</sup> | 84 | 681 | 3,223 |
| biofilm | Zhang et al., 2019<sup>35</sup> | 130 | 2,577 | 3,209 |
| GEOTRACES | Biller et al., 2018<sup>18</sup> | 610 | 4,998 | 3,063 |
| North Sea | Kruger et al., 2019<sup>56</sup> | 38 | 832 | 3,019 |
| *Tara* polar | Salazar et al., 2019<sup>24</sup> | 41 | 1,416 | 2,762 |
| *Tara* girus | Sunagawa et al., 2015<sup>22</sup> | 59 | 1,612 | 2,757 |
| Baltic Sea | Alneberg et al., 2018<sup>28</sup> | 81 | 566 | 2,335 |
| Mediterranean | Lopez-Perez et al., 2017<sup>71</sup>,<br>Haro-Moreno et al., 2019<sup>72</sup>,<br>Martin-Cuadrado et al., 2015<sup>73</sup> | 37 | 599 | 2,292 |
| HOT | Mende et al., 2017<sup>30</sup> | 85 | 1,000 | 2,109 |
| Malaspina | Acinas et al., 2021<sup>25</sup>,<br>Gregory et al., 2019<sup>23</sup> | 72 | 209 | 1,320 |
| Med. coastal | Galand et al., 2018<sup>27</sup> | 40 | 276 | 1,243 |
| Canada Basin | Colatriano et al., 2018<sup>19</sup> | 12 | 362 | 1,083 |
| Hawaii bloom | Wilson et al., 2017<sup>74</sup> | 88 | 530 | 641 |
| San Pedro Channel | Sieradzki et al., 2019<sup>31</sup>,<br>Ignacio-Espinoza et al., 2019<sup>75</sup> | 65 | 1,527 | 554 |
| sediment trap | Poff et al., 2021<sup>34</sup> | 63 | 470 | 506 |
| low oxygen | Thrash et al., 2017<sup>6</sup>,<br>Tsementzi et al., 2016<sup>76</sup>,<br>Glass et al., 2015<sup>77</sup> | 26 | 123 | 476 |
| Atlantic | Bergauer et al., 2018<sup>78</sup> | 7 | 180 | 451 |
| Red Sea | Haroon et al., 2016<sup>79</sup> | 45 | 83 | 319 |
| NW Pacific | Saw et al., 2020<sup>10</sup>, Li et al., 2018<sup>80</sup> | 35 | 96 | 248 |
| Baltic Sea virus | Nilsson et al., 2019<sup>81</sup> | 25 | 261 | 222 |

### Sequence assemblies and metagenome binning

Metagenomic sequence data in a paired-end layout was downloaded from NCBI SRA and quality controlled by using Trimmomatic^36^ v0.35, with an option ‘LEADING:20 TRAILING:20 MINLEN:60’. If one side of the pair was discarded due to its low quality, the other was retained when it passed the quality control. The qualified reads were assembled in a sample-by-sample manner (i.e., all qualified reads from one sample were used in one assembly; statistics are described in Supplementary File S1) using MEGAHIT ^37^ v1.1.4. Resulting contigs were retained if the length is no less than 1 kbps.

We then calculated a coverage profile for each metagenome using all metagenomes belong to the same division for better binning performance (Table 1; see also ‘Technical Validation’). An exception was applied to the division of GEOTRACES, which includes many metagenomes (n=610). This division is split into six subdivisions and the coverage profiles were calculated within each subdivision (Supplementary File S1). Read mapping was performed by bowtie2^38^ v2.3.5.1 using qualified paired-end reads. Mapping result was sorted by samtools (http://www.htslib.org/) v1.9 and coverage was calculated by jgi_summarize_bam_contig_depths that is bundled in MetaBAT2^39^, customizing a parameter ‘-percentIdentity’ set to 90. We then performed metagenome binning using three algorithms, MetaBAT2^39^ v2.12.1, MaxBin2^40^ v2.2.6, and CONCOCT^41^ v1.0.0. These algorithms were run with default settings, but for MetaBAT2, the ‘--minContig’ parameter was set to 1,500 following the software instruction, which states this value should not be less than 1,500. The resulting three sets of bins were then dereplicated and merged using the bin_refinement module of MetaWRAP^42^ v1.2.1 with minimum completion is set to 50. The quality score (QS) was defined as ‘%-completeness - 5 x %-contamination’ and genomes of QS ≥ 50 were retained. Completeness and contamination of genome bins was estimated by taxon specific sets of single-copy marker genes through the lineage-specific workflow of CheckM v1.0.13^43^. After removal of genomes likely derived from internal standard (n=63; *Thermus thermophilus* and *Blautia producta*^44^), 54,614 genomes were obtained.

### Post-refinement of genome bins

For quality improvement of the reconstructed genome bins, we developed a post-refinement module to decontaminate potential misassigned contigs for each genome bin (Fig 1c; see also ‘Technical Validation’). This module consists of three independent decontamination filters: (1) taxonomic filter, (2) mobile element filter, and (3) outlier filter. First, the taxonomic filter was designed to detect taxonomically inconsistent contigs with each genome. Coding regions were predicted with prodigal^45^ v2.6.3 and resulting proteins were used as input of CAT and BAT^46^ v5.0.3 to assign taxonomy for contigs and genomes, respectively. CAT and BAT was run with the default setting using NCBI Taxonomy downloaded in January 2020. Then, predicted taxonomy was quality controlled to remove less reliable assignment. Namely, predicted taxonomy was recursively trimmed from the low level until either of the following three types of assignment are not detected: (a) ‘suggestive’ taxonomic assignment that is less confident, indicated by stars in the BAT and CAT output, (b) very low-level assignment equal to or lower than species-level, and (c) some ambiguous assignment (i.e., classified as ‘environmental samples’ or classifications start with ‘unclassified’). For each pair of a genome and its contig, the pair was recognized as taxonomically consistent only if the lowest common ancestor of the genome and the contig was the same as either of these. For example, suppose taxonomy of a genome is ‘A; B; C’, a contig is taxonomically consistent if taxonomy of a contig is ‘A; B’ or ‘A; B; C; D’, and inconsistent if ‘A; B; E’ or ‘A; F’.

Second, the mobile element filter was designed to remove possible contamination of viral and plasmid contigs within genome bins. As genome bins are likely contaminated with viral and plasmid contigs that have similar coverage and nucleotide composition to the genome^20^, we adopted a conservative approach to remove possible mobile elements, though these contigs are possibly true parts of the genome (e.g., as a provirus). First, circular contigs were identified as potential viral and plasmid contigs by detecting terminal redundancy through ccfind (https://github.com/yosuken/ccfind)^47^. Second, viral contigs were detected using additional two types of methods. VirSorter^48^ v1.0.6 was used to detect viral contigs of ≥3kb. The prediction result of category 1-6 was considered as viral, but for category 4-6 (predicted as provirus), only if length of viral region was ≥50% of the total length, the contig was considered as viral. To supplement the detective power for short contigs (1kb to 10kb), we additionally scanned for *terL* genes that are one of the hallmark genes of prokaryotic viruses, by following steps. We prepared 11 *terL* HMMs (Supplementary File S2) that were constructed from *terL* protein sequences obtained from previously identified aquatic viral MAGs (EVGs: circularly assembled environmental viral genomes)^47^. We searched for *terL* candidates using hmmsearch (HMMER^49^ v3.2.1; evalue < 1e-10) with the 11 HMMs as query. We validated sequence homology of the candidates with known *terL* genes using pipeline_for_high_sensitive_domain_search (https://github.com/yosuken/pipeline_for_high_sensitive_domain_search), which utilizes jackhmmer (HMMER^49^ v3.2.1) to build a protein HMM of each gene and hhsearch^50^ (HH-suite^50^ v3.2.0) to identify homology between the built HMMs and *terL* HMMs included in pfam 32.0. The candidates were identified as *terL* if the best hit is one of *terL* domains (i.e., Terminase_1, Terminase_3, Terminase_6, Terminase_GpA, DNA_pack_N, Terminase_3C, and Terminase_6C) among all the pfam domain and if probability of the HHsearch hit is >97%. We used proteins encoded in EVGs as a database of jackhmmer (jackhmmer parameters: ‘-N 5 --incE 0.001 --incdomE 0.001’).

Third, the outlier filter was designed to detect outlier contigs in terms of coverage and tetranucleotide frequency (<-2.5 or >2.5 s.d. within each genome bin). Principal component analysis was performed using the prcomp function of R v3.6.2 (with default parameters) and the first primary component was evaluated. As a coverage profile, a part (related to contigs of the bin) of a coverage profile that was used for binning was extracted and normalized within each sample. Contigs identified as outliers were removed from the genome bin. Overall, after the detection and removal of possible contamination using these three filters, completeness and contamination of each genome bin was again estimated with the lineage-specific workflow of CheckM.

Finally, 52,325 genomes of QS ≥ 50 were obtained and here named as the OceanDNA MAGs (Data Citation 1; Table S2). The OceanDNA MAGs reconstructed from various marine environments and size-fractions (Fig 2a), including deep-sea deeper than 1,000m (3,337 genomes from 176 samples), low-oxygen zones of <90 μmol O_2_ per kg water (7,884 genomes from 176 samples), polar regions (7,752 genomes from 129 samples), viral enriched fraction (pass through a filter of 0.2 or 0.22 μm pore; 5,998 genomes from 312 samples). Basic statistics of genome assemblies were evaluated with QUAST^51^ v5.0.2 (Supplementary File S3). Ribosomal RNAs and transfer RNAs were identified using Barrnap v0.9 (https://github.com/tseemann/Barrnap) and tRNAscan-SE^52^ v2.0.5, respectively.

### Taxonomic assignment and their novelty evaluation using GTDB

We performed species-level clustering and identified species representatives of the OceanDNA MAGs through the following two rounds. First, for each of the 24 divisions, species-level clustering was performed using dRep^53^ v2.2.2 with a cutoff value of average nucleotide identity set as 95% and aligned fraction as 30%. We identified genomes of species representatives if ‘QS + ln(N50)’ was the highest within each species-level cluster. From the 24 divisions, 13,357 species representatives were identified at this round. Then, the secondary clustering was performed among these representatives using dRep, and 8,466 species-level clusters were obtained. The representatives of the 8,466 species-level clusters (Data Citation 2) were identified using the same criteria. The median genome completeness and contamination of both the species representatives and the other genomes (n=43,859; Data Citation 3) were estimated as >80% and <2%, respectively (Fig 2b). The species representatives were originated from various metagenomic projects and not dominated by ones from *Tara* Oceans (Fig. 2c).

The OceanDNA MAGs were taxonomically classified using GTDB (Genome Taxonomy DataBase) release 05-RS95^21^ through the classify workflow of GTDB-Tk^54^ v1.3.0. As classification based on GTDB, the species representatives spanned 59 phyla (Fig. 2d). Of these, 11 species representatives were not assigned to any existing class, suggesting that these species potentially belong to new classes. Likewise, it was suggested that 44 species representatives belong to new orders, 290 belong to new families, and 1,395 belong to new genera and 4,516 belong to new species (Fig. 2e). Overall, most of the species representatives (n=6,256; 73.9%) were not assigned to existing species in the database.

### Novelty evaluation using published marine genomes

For further novelty assessment of the OceanDNA MAGs, we comprehensively collected published genomes of marine prokaryotes. First, genomes contained in MarDB and MarRef^55^ v5.0, which are curated genome collection of marine prokaryotes originated from isolates/SAGs/MAGs, were downloaded (n=14,209). Second, to supplement these with very recently published genomes and/or genomes that are not stored in NCBI, we collected genomes (n=26,946; SAGs and MAGs) of marine origin from 15 research articles^3,5,6,10,24,33,35,56,57,58,59,60,61,62,63^ (Supplementary File S4). After selection of qualified genomes (QS ≥ 50), 29,292 genomes were retained in total (11,985 from marRef/MarDB and 17,307 genomes from the 15 articles; Supplementary File S5). We then organized a unified genome catalog of marine prokaryotes (UGCMP; n=81,617), composed of the 29,292 published genomes and the 52,325 OceanDNA MAGs (Fig. 2f). We identified species representatives of UGCMP by following two steps. Species-level clusters (n=13,669) and the representatives were identified separately for MarDB/MarRef and for each publication, using the same criteria as the OceanDNA MAGs. After unifying the species representatives of OceanDNA MAGs (n=8,466) and published marine genomes (n=13,669) into one set, the second-round species-level clustering was performed with the same conditions. We finally identified 16,141 species representatives of UGCMP using the same criteria (Supplementary File S6). The OceanDNA MAGs exclusively composed 4,806 species-level clusters (56.8% of the species representatives of the OceanDNA MAGs) and selected as species representatives in 1,971 non-exclusive species-level clusters (23.3% of the species representatives of OceanDNA MAGs) based on the better genome quality evaluated by ‘QS + ln(N50)’. Overall, a large part (80.1%; n=6,777) of the species representatives of the OceanDNA MAGs was still species representatives in UGCMP.

We then assessed phylogenomic diversity of UGCMP for bacteria (n=74,214) and archaea (n=7,403). For domain and phylum-level classification, taxonomic assignment of UGCMP genomes were performed using GTDB release 05-RS95 and GTDB-Tk v1.3. Phylogenomic trees of bacteria and archaea were reconstructed with FastTree v2.1.11 (option: ‘-wag -gamma’) using alignments that were built by GTDB-Tk (Fig 2g). The alignments included 5,040 sites of high phylogenetic signal from 120 single copy marker genes for bacteria, and 5,124 sites from 122 genes for archaea as well. After midpoint rooting using gotree (https://github.com/evolbioinfo/gotree) v0.4.0, sum of branch length was calculated for two categories: (1) branches that were represented only by the OceanDNA MAGs (2) branches that were other than (1). The expanded phylogenetic diversity by the OceanDNA MAGs was 34.2% (34.8% for bacteria and 29.4% for archaea), estimated from a ratio of (1) to (2).

### Back mapping of metagenomic reads

We assessed the fraction of metagenomic reads recruited onto the OceanDNA MAGs. Sequence reads of the 2,057 metagenomes, which were used for genome reconstruction, were back mapped onto the 8,466 species representatives of the OceanDNA MAGs. For cases that one sample has multiple sequencing runs, only the biggest run was used. Read mapping was performed with bowtie2 ^38^ v2.3.5.1 with the default setting using the quality controlled paired-end reads of each run, but if the run was bigger than 5 Gbps, a subset of 5 Gbps sequences that was randomly sampled using seqtk (https://github.com/lh3/seqtk) v1.3 was used for read mapping. Then, the mapping result was sorted using samtools (http://www.htslib.org/) v1.9, and only mapping of ≥95% identity, ≥80 bp, and ≥80% aligned fraction of the read length was extracted using msamtools (https://github.com/arumugamlab/msamtools) that are bundled in MOCAT2^64^ v2.1.3. Finally, the mapped reads were counted using featureCounts^65^ that were bundled in Subread v2.0.0. The species representatives collectively cover 10.4 - 35.0% (the 25th to 75th percentile) of metagenome reads of the 2,057 metagenomes (Fig 3a). Especially, where only prokaryotes-enriched metagenomes (n=731) were considered, 26.5 - 42.0% of metagenomic reads were mapped onto the species representatives.

Next, we evaluated mapped read fractions onto species representatives of UGCMP, the OceanDNA MAGs, and four sets of marine prokaryotic genomes from large-scale genome reconstruction studies^3,5,16, 62^ (Fig 3b). Read mapping was performed using only species representatives of qualified genomes (i.e., QS ≥ 50) for all these genome collections. In terms of the medians of mapped read fractions, the OceanDNA MAGs was the highest among the previously reported genome collections, and UGCMP was about 10% higher than the OceanDNA MAGs.

## Data Records

Genome sequences of the OceanDNA MAGs (Data Citation 1) were available at figshare (https://figshare.com/s/e2aa3456d68aa51e617c) and submitted to DDBJ/ENA/GenBank under BioProject accession no. PRJDB11811. Genome sequences of the 8,466 species representatives (Data Citation 2) were submitted as WGS entries, and sequences of the other genomes (n=43,859) were submitted as DDBJ analysis entries (Data Citation 3; available via DDBJ). Supplementary files (listed below) are available at figshare (https://figshare.com/s/e2aa3456d68aa51e617c). Related information of the OceanDNA MAGs can be accessed at https://OceanDNA-MAGs.aori.u-tokyo.ac.jp.

**Supplementary File S1.** A list of metagenomes used in this study, with various information used for generating figures.

**Supplementary File S2.** Eleven multiple alignments and HMMs of *terL* protein sequences obtained from aquatic viral MAGs.

**Supplementary File S3.** A list of the OceanDNA MAGs with basic statistics, functional RNAs, genome quality, and genome-based taxonomy.

**Supplementary File S4.** A list of 15 publications of marine SAGs and MAGs

**Supplementary File S5.** A custom collection of published marine prokaryotic genomes of QS ≥ 50

**Supplementary File S6.** A list of species representatives of UGCMP

## Technical Validation

For maximization of the genome quality, our genome reconstruction pipeline was carefully designed, including three key processes (Fig. 1c): (1) high-resolution coverage profiles were calculated using all metagenomes within each division, (2) metagenome binning was performed using three algorithms and subsequently dereplicated, (3) an automated post-refinement process for detection of possible contaminations, including ones could be missed by prokaryotic single-copy marker gene-based assessment. Here we assessed efficiency of these processes.

First, binning algorithms are primarily based on a coverage profile among multiple metagenomes and *k*-mer (e.g., tetranucleotide) composition of metagenomic contigs^66,67^. It was shown that if a coverage profile was calculated using only a few metagenomes, it would result in underperformance of a binning algorithm (e.g., CONCOCT)^41^. Here, to assess the effect of the number of metagenomes in coverage profile, we selected 20 *Tara* Oceans metagenomes included in the “*Tara* prok” division (Table 1) of which geographic region and water depth was widely distributed. We performed metagenome binning of the selected metagenomes with different coverage profiles. The coverage profiles were calculated with all metagenomes within the same division (n=139) or with randomly sampled 10, 25, and 50 metagenomes with three replicates out of the 139 metagenomes. If multiple sequencing runs were available from one metagenome, only the largest run was used for coverage profiles. Then, binning was performed same as the OceanDNA MAGs but except for the post-refinement part, and the resulting number of bins of QS ≥ 50 was compared (Fig 4a). As a result, coverage profiles of all metagenomes reconstructed the greater number of qualified bins (i.e., QS ≥ 50) than coverage profiles of subsampled metagenomes. The result suggests the superiority of the ‘high-resolution’ coverage profiles calculated with many metagenomes.

**Figure 4.**
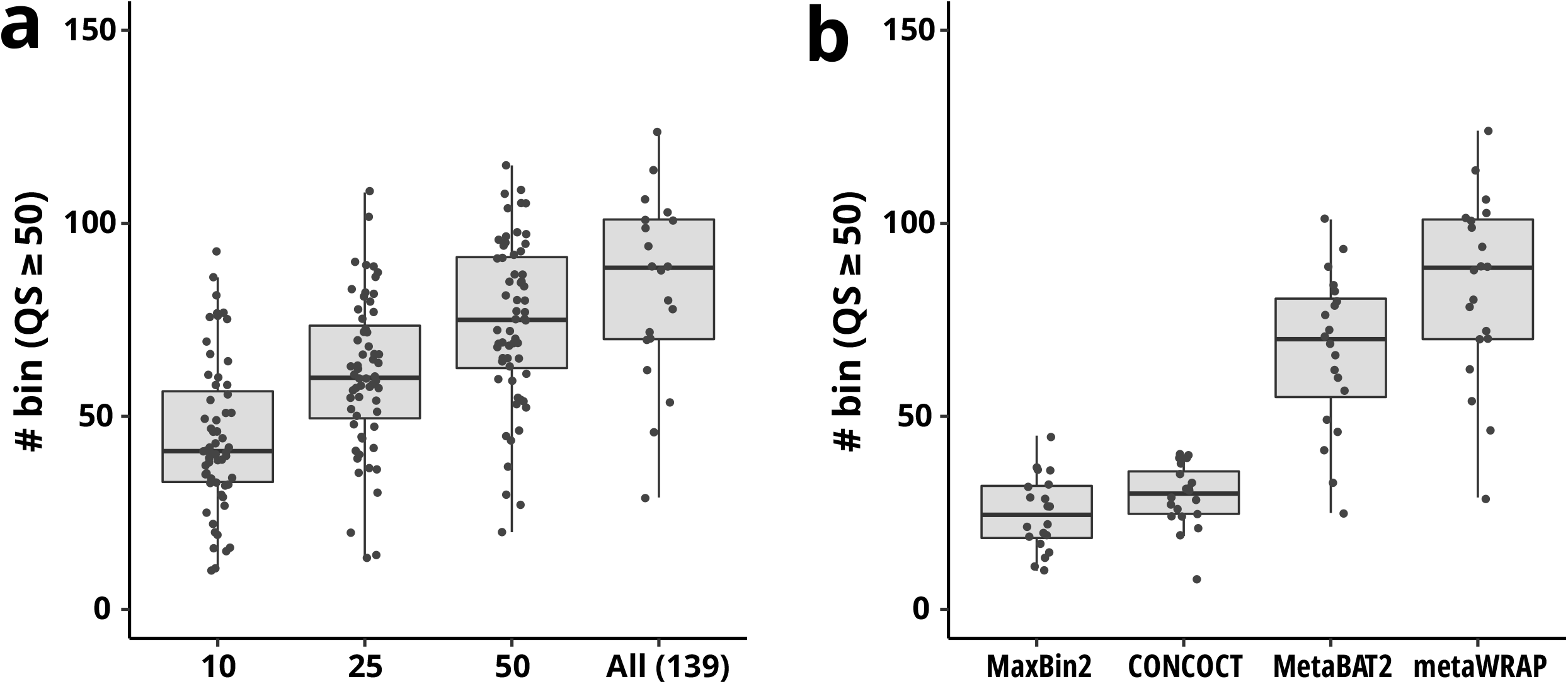
Assessment of the genome reconstruction pipeline. Using selected 20 *Tara* Oceans metagenomes included in the “*Tara* prok” division, the impact of high-resolution coverage profiles (a) and use of multiple binning algorithms (b) were assessed. The number of qualified genome bins (QS ≥ 50) was compared between (a) coverage profiles calculated with all metagenomes within the same division (n=139) or with randomly sampled 10, 25, and 50 metagenomes with three replicates, and (b) different algorithms: MaxBin2, CONCOCT, MetaBAT2, and dereplicated results of the three algorithms using the bin_refinement module of MetaWRAP.

Second, using the same 20 metagenomes of the “*Tara* prok” division, binning result of single algorithm (MetaBAT2, CONCOCT, MaxBin2) and dereplicated result of the three algorithms using the bin_refinement module of MetaWRAP were compared (Fig 4b). Dereplication of bins generated from three algorithms significantly increased the number of qualified (i.e., QS ≥ 50) bins.

Third, we designed an automated post-refinement process to remove possible contamination from each MAG using three filters that are independent of prokaryotic single-copy marker genes: (1) taxonomic filter, (2) mobile element filter, and (3) outlier filter. Similar strategies were applied in previous studies (e.g., MAGpurify^68^). The aim of this refinement process is to remove possible contamination for genome quality improvement. Especially, contamination over the domain (i.e., eukaryotic and viral contigs included in prokaryotic genomes) could not be detected through analysis of prokaryotic single-copy marker genes. Genomes from *Tara* Oceans MAG studies were predicted to contain viral contigs (in a few cases, more than 50) within a single genome^69^. Viral contigs could be contamination of viral genome fragments that have similar coverage profiles and k-mer compositions to the prokaryotic genome^20^. Though removing viral and plasmid sequences possibly results in the exclusion of true element of the genome (e.g., provirus) and identification of viral and plasmid contigs could contain false positives, we set a priority on removing those as possible contamination, not retaining those as true genomic fragments. The three filters of the post-refinement module identified 561,804, 39,289, and 436,143 potential misassigned contigs, respectively. Overall, from 54,614 qualified genome bins, 1,000,417 contigs were filtered out (18.3 contigs per genome bin on average) and 2,289 genome bins were discarded, due to the reduction of genome completeness as a result of the decontamination process. Code for the post-refinement process is available at GitHub (https://github.com/yosuken/MAGRE).

## Usage Notes

We carefully designed the genome reconstruction pipeline for genome quality improvement, including the automated post-refinement process. Nevertheless, due to difficulty of perfect decontamination, misassigned contigs might be still included in the genomes. Manual quality control is recommended before use of the genomes, as is the case for MAGs from other studies.

We collected metagenome samples covering various marine environments. Nevertheless, note that some marine environments (e.g., hydrothermal vents, sediments, coral reefs, and oil spills) were not included in the dataset of this study.

Genome completeness evaluated by CheckM are likely underestimated for genomes of specific taxa that have experienced extreme genome reduction and may have a symbiotic lifestyle (e.g., lineages of the phylum Patescibacteria, also known as Candidate Phyla Radiation). Ribosomal RNA operons are difficult regions to reconstruct due to co-existence of closely related sequences that confuse de Bruijn graph-based assemblers^20^. 5S, 16S, 23S ribosomal RNAs were identified in 24.2%, 6.8%, 3.8% of the OceanDNA MAGs, respectively (including full sequences or >25% fragments of the whole length).

SAR11 and Prochlorococcus are two of the most abundant lineages in the ocean. However, despite their high abundance, not so many genomes of these lineages were reconstructed in this study. This shortfall is probably attributable to coexisting closely related strains of these lineages that cause difficulty for genome reconstruction^20^. Among the OceanDNA MAGs, 780 genomes were reconstructed from 85 species-level clusters of ‘o__Pelagibacterales’ (SAR11) and 157 genomes were reconstructed from 8 species-level clusters of ‘g__Prochlorococcus’. For these lineages, SAGs could supplement genomic information. For example, recently reported SAGs that were reconstructed from the tropical and subtropical euphotic ocean^5^ includes 2,108 genomes consisted of 1,215 species-level clusters of ‘o__Pelagibacterales’ and 327 genomes consisted of 155 species-level clusters of ‘g__Prochlorococcus’, where genomes are limited to those of QS ≥ 50 (Supplementary File S5).

## Code Availability

Code of the post-refinement module is available at GitHub as MAGRE (https://github.com/yosuken/MAGRE).

The options and parameters of all tools used for the analysis are described in the main text.

## Acknowledgements

We thank all persons who contributed to generation of the metagenome sequence data, as well as all persons who developed the software and databases used in this study. This study is partly supported by JSPS KAKENHI Grant Number 18K19224, 18H04136, and 21K19134 (S.Y.). Computation time was provided by the Super Computer System, Institute for Chemical Research, Kyoto University.

## Author contributions

Y.N. conceived the study, designed and implemented the pipeline, performed analysis, and wrote a draft. S.Y. reviewed and edited a draft.

